# Annotation of Putative Circadian Rhythm-Associated Genes in *Diaphorina citri* (Hemiptera : Liviidae)

**DOI:** 10.1101/2021.10.09.463768

**Authors:** Max Reynolds, Lucas de Oliveira, Chad Vosburg, Thomson Paris, Crissy Massimino, Jordan Norus, Yasmin Ortiz, Michelle Espino, Nina Davis, Ron Masse, Alan Neiman, Rachel Holcomb, Kylie Gervais, Melissa Kemp, Maria Hoang, Teresa D. Shippy, Prashant S. Hosmani, Mirella Flores-Gonzalez, Lukas A. Mueller, Wayne B. Hunter, Joshua B. Benoit, Susan J. Brown, Tom D’Elia, Surya Saha

## Abstract

The circadian rhythm is a process involving multiple genes that generates an internal molecular clock, allowing organisms to anticipate environmental conditions produced by the earth’s rotation on its axis. This report presents the results of the manual curation of twenty-seven genes likely associated with circadian rhythm in the genome of *Diaphorina citri*, the Asian citrus psyllid. This insect acts as the vector of the bacterial pathogen *Candidatus* Liberibacter asiaticus (CLas), the causal agent of citrus greening disease (Huanglongbing). This disease is the most severe detriment to citrus industries and has drastically decreased crop yields worldwide. Based on the genes identified in the psyllid genome, namely *cry1* and *cry2, D. citri* likely possesses a circadian model similar to that of the lepidopteran butterfly, *Danaus plexippus*. Manual annotation of these genes will allow future molecular therapeutics to be developed that can disrupt the psyllid biology.

## Introduction

Huanglongbing (HLB), or citrus greening disease, is caused by the bacterium *Candidatus* Liberibacter asiaticus (CLas) infecting citrus phloem, causing loss of fruit production and eventually tree death [1,2]. The Asian citrus psyllid (*Diaphorina citri*) acts as an effective vector for the pathogen, thereby spreading the disease. Currently there is no effective treatment to stop the disease, and it has caused widespread crop damage, resulting in substantial financial losses in the citrus industry [3]. To prevent further damages, development of management strategies, such as molecular therapeutics, are being investigated which requires a solid foundation of the genetic-basis of psyllid biology. To better understand *D. citri* biology, we first need to identify gene pathways within the psyllid genome. To this end, genes involved in the main circadian rhythm pathway loops, along with ancillary circadian rhythm genes, were manually annotated in the *D. citri* genome (Table 2). Based on the critical importance of the genes identified, many are promising targets for development of molecular therapeutics based on RNAi strategies in insects (Table 1). Disruption of *D. citri* circadian rhythm may alter psyllid behavior, potentially hindering the spread of HLB.

**Table 1:** List of annotated genes related to circadian rhythm with their corresponding RNAi studies and references.

| Genes | Organism | RNAi outcome | Reference |
| --- | --- | --- | --- |
| ***cycle* and *Mip*** | *Plautia stali* | RNAi of *cyc* in conjunction with *myoinhibitory peptide (Mip)* induced diapause by disrupting photo periodic response in ovarian development | Tamai T, Shiga S, Goto SG. Roles of the circadian clock and ENDOCRINE regulator in THE photoperiodic response of the Brown-winged green Bug *Plautia stali*. Physiological Entomology. 2018;44(1):43–52. doi:10.1111/phen.12274 |
| ***cycle*** | *Riptortus pedestris* | diapause was induced under a diapause-averting photoperiod | Ikeno T, Tanaka SI, Numata H, Goto SG. Photoperiodic diapause under the control of circadian clock genes in an insect. BMC Biology. 2010;8(1):116. doi:10.1186/1741-7007-8-116 |
| **period** | *Riptortus pedestris* | diapause was prevented under a diapause-inducing photoperiod | Ikeno T, Tanaka SI, Numata H, Goto SG. Photoperiodic diapause under the control of circadian clock genes in an insect. BMC Biology. 2010;8(1):116. doi:10.1186/1741-7007-8-116 |
| **period** | *Ostrinia nubilalis* | Underlie a polymorphism among populations that result in the early termination of diapause | Kozak GM, Wadsworth CB, Kahne SC, Bogdanowicz SM, Harrison RG, Coates BS, Dopman EB. Genomic basis of CIRCANNUAL rhythm in the European Corn Borer Moth. Current Biology. 2019;29(20). doi:10.1016/j.cub.2019.08.053 |
| **Pigment-dispersing factor receptor** | *Ostrinia nubilalis* | Affects diapause occurrence and may influence seasonal timing through alteration of circadian clock function | Kozak GM, Wadsworth CB, Kahne SC, Bogdanowicz SM, Harrison RG, Coates BS, Dopman EB. Genomic basis Of CIRCANNUAL rhythm in the European Corn Borer Moth. Current Biology. 2019;29(20). doi:10.1016/j.cub.2019.08.053 |
| **Clock** | *Schistocerca gregaria* | Knockdown of *clk* caused mortality in female insects before breeding age | Tobback, J., Boerjan, B., Vandersmissen, H.P. and Huybrechts, R. (2011) The circadian clock genes affect reproductive capacity in the desert locust *Schistocerca gregaria*. Insect Biochem Mol Biol 2011;41(5):313–321. doi:10.1016/j.ibmb.2011.01.008 |
| **period** | *Schistocerca gregaria* | *per* and *tim* RNAi resulted in reduced progeny suggesting that PER and TIM are needed in the ovaries for egg development | Tobback, J., Boerjan, B., Vandersmissen, H.P. and Huybrechts, R. (2011) The circadian clock genes affect reproductive capacity in the desert locust *Schistocerca gregaria*. Insect Biochem Mol Biol 2011;41(5):313–321. doi:10.1016/j.ibmb.2011.01.008 |
| **timeless** | *Schistocerca gregaria* | *tim* knockdown affected female insect egg deposition and resulted in reduced offspring | Tobback, J., Boerjan, B., Vandersmissen, H.P. and Huybrechts, R. (2011) The circadian clock genes affect reproductive capacity in the desert locust *Schistocerca gregaria*. Insect Biochem Mol Biol 2011;41(5):313–321. doi:10.1016/j.ibmb.2011.01.008 |
| **vrille** | *D. melanogaster* | Dominant maternal enhancer of defects in dorsoventral patterning causing embryo mortality. | George H, Terracol R. 1997. The *vrille* gene of *Drosophila* is a maternal enhancer of *decapentaplegic* and encodes a new member of the *bZip* family of transcription factors. Genetics. 146:1345-1363 |
| **takeout** | *D. melanogaster* | Mutant *takeout* causes abnormal locomotor activity resulting in insect death from starvation. | Sarov-Blat L, Venus So W, Liu L, Rosbash M. 2000. The *Drosophila Takeout* gene is a novel molecular link between circadian rhythms and feeding behavior. Cell. 101: 647-656 |

**Table 2.** Evidence supporting gene annotation. There are twenty-seven annotated circadian rhythm genes in *D. citri*. Each gene model has been assigned an Official Gene Set (OGS) version 3.0 gene identifier, and the evidence types used to validate or modify the structure of the gene model have been listed. Genes marked as complete have a complete coding region and those not marked as complete were only able to be annotated as partial gene models. Descriptions of the various evidence sources and their strengths and weaknesses are included in the online protocol [28]. Terminology is as in flybase.org [30].

| Gene | OGSv3 ID | MCOT ID | de novo | Gene Model | Evidence Supporting annotation |  |  |  |
| --- | --- | --- | --- | --- | --- | --- | --- | --- |
|  |  |  |  | Complete / Partial | MCOT | Iso-Seq | RNA-Seq | Ortholog |
| *Drosophila* Model Core Clock Genes |  |  |  |  |  |  |  |  |
| *circadian locomotor output cycles protein kaput* | Dcitr05g05480.2.1 | MCOT03010.0.CO | X |  | X |  | X |  |
| *cycle* | Dcitr05g16440.2.3 |  |  | X |  |  | X | X |
| *PAR domain protein 1* | Dcitr05g02780.2.1 | MCOT14609.2.CT |  | X | X |  | X |  |
| *period* | Dcitr09g03540.1.1 | MCOT12213.0.CO |  | X | X |  | X | X |
| *timeless* | Dcitr04g02940.1.1 | MCOT23350.0.CT | X | X | X | X | X | X |
| *vrille* | Dcitr10g05230.1.1 | MCOT16283.2.CT | X | X | X | X |  |  |
| **Other Negative Loop Associated Genes** |  |  |  |  |  |  |  |  |
| *Casein kinase 2 alpha subunit* | Dcitr02g18000.1.1 | MCOT21500.0.CO | X | X | X |  | X | X |
| *Casein kinase 2 beta subunit* | Dcitr11g02180.1.1 |  | X | X |  | X |  | X |
| *Cryptochrome 1* | Dcitr07g01550.1.1 | MCOT00625.0.CT | X | X | X | X | X | X |
| *Cryptochrome 2* | Dcitr09g09100.1.1 | MCOT19207.3.CO | X | X | X |  | X |  |
| *double-time* | Dcitr07g07760.2.1 | MCOT08150.0.MT |  | X | X |  | X |  |
| **Other Positive Loop Associated Genes** |  |  |  |  |  |  |  |  |
| *ecdysone-induced protein 75* | Dcitr12g07770.2.1 | MCOT02379.0.CT |  | X | X | X | X | X |
| *Hormone Receptor 3 /Rora* | Dcitr08g04060.1.1 | MCOT23147.0.CC |  | X | X | X |  |  |

| Genes Documented to be Under Circadian Influence |  |  |  |  |  |  |  |  |
| --- | --- | --- | --- | --- | --- | --- | --- | --- |
| *clockwork orange isoform 1* | Dcitr01g06780.1.1 | MCOT17066.1.CO |  | X | X |  | X |  |
| *clockwork orange isoform 2* | Dcitr01g06780.1.1 | MCOT17066.2.CO | X | X | X |  | X |  |
| *Ecdysone receptor* | Dcitr02g19530.1.1 | MCOT15763.0.CT |  | X | X | X | X | X |
| *Hormone Receptor 3 Homolog* | Dcitr08g02450.1.3 |  | X | X |  |  |  | X |
| *insulin related peptide* | Dcitr06g08750.1.1 | MCOT08282.0.MO |  | X | X |  | X |  |
| *lark* | Dcitr01g16940.1.1 | MCOT03363.0.CT | X | X | X | X | X | X |
| *ovary maturing parsin* | Dcitr09g03370.1.1 | MCOT04488.1.CT |  | X | X |  |  |  |
| *Protein phosphatase PP2A 55 kDa regulatory subunit* | Dcitr09g05720.1.1 | MCOT08076.0.OO | X | X | X | X | X | X |
| *Serine/Threonine-Protein phosphatase 2A 56 kDa regulatory subunit epsilon* | Dcitr03g10950.1.1 | MCOT02984.0.CT | X | X | X | X | X | X |
| *Serine/Threonine-Protein phosphatase 2A 56 kDa regulatory subunit epsilon* | Dcitr03g03510.1.1 | MCOT08848.0.MO | X | X | X |  | X | X |
| *takeout 1* | Dcitr07g054<br>80.1.3 | MCOT2100<br>1.2.CT | X | X | X | X | X | X |
| *takeout 2* | Dcitr06g072<br>20.1.1 | MCOT1641<br>3.0.CO |  | X | X | X | X | X |
| *takeout-Like* | Dcitr09g076<br>20.1.1 | MCOT1335<br>7.0.CC | X | X | X | X | X | X |
| *timeless 2/*<br>*timeout*<br>*fragment 1* | Dcitr00g113<br>40.2.1 |  |  |  |  |  | X | X |
| *timeless 2/*<br>*timeout*<br>*fragment 2* | Dcitr00g055<br>20.2.1 |  |  |  |  |  |  | X |
| *timeless 2/*<br>*timeout*<br>*Fragment*<br>*3* | Dcitr00g055<br>10.2.1 |  |  |  |  |  | X | X |
| *vestigial* | Dcitr08g079<br>20.1.1 | MCOT0681<br>0.0.CC | X | X | X |  | X | X |
| *Vitellogeni*<br>*n-1* | Dcitr08g093<br>20.1.1 |  |  | X |  | X |  | X |
| *Vitellogeni*<br>*n-2* | Dcitr01g056<br>60.1.1 | MCOT0528<br>2.2.CO |  | X | X | X | X |  |

## Context

The circadian rhythm is a process regulated by the interactions between multiple genes, allowing a cell to respond to, and anticipate day-night conditions during a roughly 24-hour cycle [4,5]. Periodicity occurs due to fluctuations in gene expression autoregulated by inhibitory feedback loops based on external stimuli (principally light, but also temperature, humidity, nutrition and others), also known as *Zeitgeibers* [5-9]. The basic mechanism for circadian clocks in animals is highly conserved across all taxa, and the best studied insect circadian system is that of the fruit fly, *Drosophila melanogaster* (Order: Diptera) [10]. In this model, the circadian rhythm is primarily regulated by six transcription factors: PERIOD (PER), TIMELESS (TIM), CYCLE (CYC), CLOCK (CLK), VRILLE (VRI), and PAR DOMAIN PROTEIN 1 (PDP1) [11]. The *D. melanogaster* model is an invaluable reference point in understanding the circadian rhythm, but it cannot be completely generalized to *D. citri* (Order: Hemiptera). This is because hemipterans are a more evolutionarily ancient lineage [12] and there exists differences in cryptochrome genes (*cry*) between insects [13-15].

In nature, individual cryptochrome proteins (CRY) have evolved to complete different functions. CRY1 operates as a blue light receptor, while CRY2 works as a circadian transcription repressor [13, 15]. Insects can possess one or both cryptochrome genes, and this affects how their feedback loops operate. These differences result in three different circadian rhythm models: the *Drosophila* model possesses *cry1*, the butterfly *Danaus plexippus* possess both *cry1* and *cry2*, and the honeybee *Apis mellifera* and the beetle *Tribolium castaneum* have only *cry2* (Fig. 1) [14, 15]. Cryptochrome variation gives possible insight into the evolution and function of the circadian rhythm in insects, but also makes the *D. melanogaster* model different from non-dipterans due to their possession of *cry2* [14].

**Figure 1.**
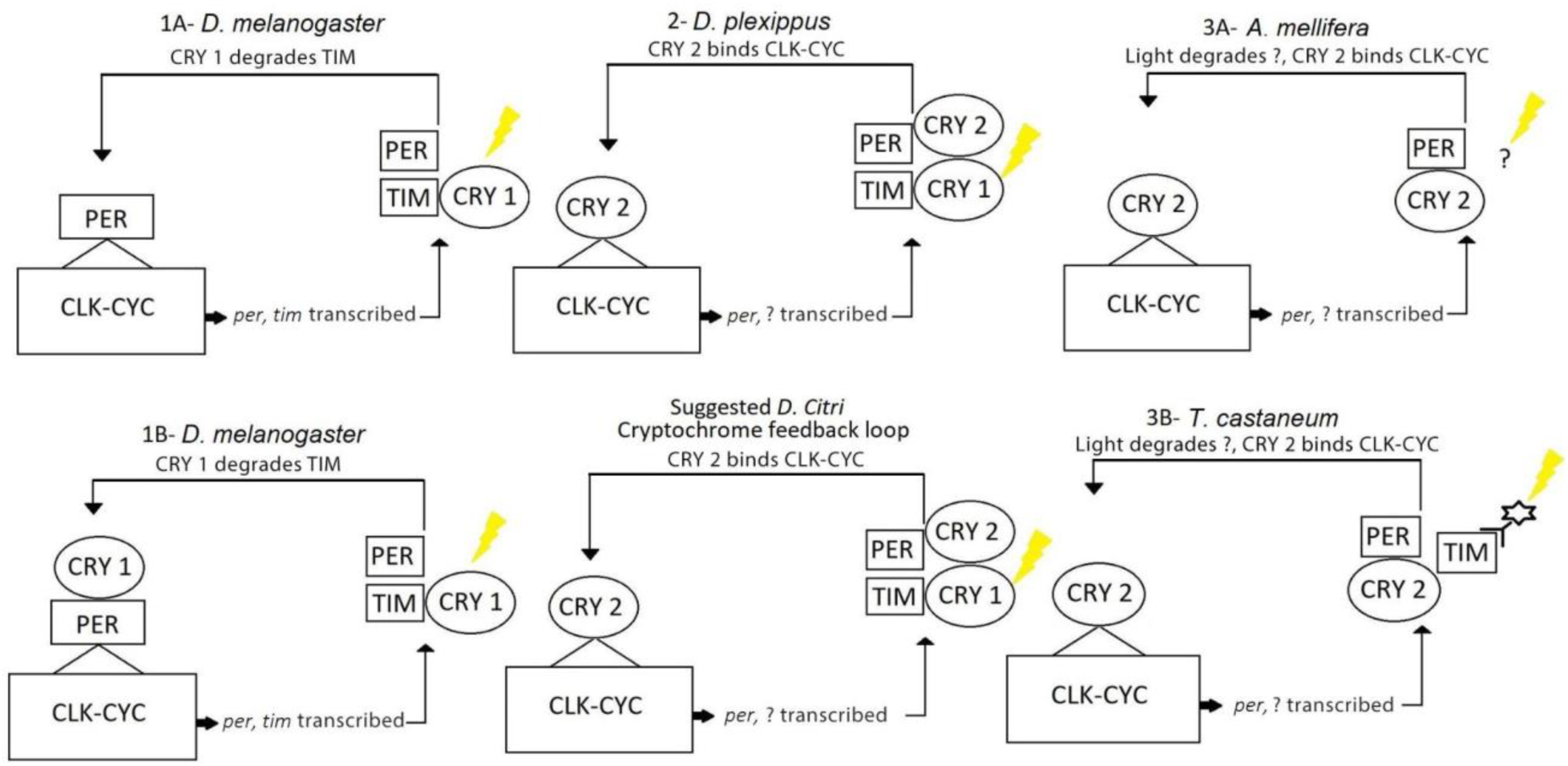
Circadian models that differ based on cryptochrome variation. *D. melanogaster* possesses only *cry1, A. mellifera* and *T. castaneum* possess only *cry2*, and *D. plexippus* possesses both. The presence of both cryptochromes in *D. citri* suggests a similar circadian model to *D. plexippus*.

In addition to cryptochrome variation between insects, the presence or absence of other genes can affect how an organism’s circadian model operates. For example, the *D. citri* genome possesses *clockwork orange (cwo), Hormone receptor 3 (Hr3)* and *ecdysone-induced protein 75 (E75)* which may suggest a *cyc* transcriptional regulation system similar to those found in *Thermobia domestica* and *Gryllus bimaculatus* [16,17].

Based on the genes identified and annotated in the *D. citri* genome, Figure 2 presents a theoretical model of the circadian rhythm pathway in *D. citri*. Regulation of the pathway operates in two feedback loops, one negative and one positive [11]. The negative loop involves the *per* and *tim* genes, whose expression is promoted when a heterodimer of CLK and CYC binds to their E-Box promoters [10, 18-21]. The CLK-CYC heterodimer is also responsible for the transcription activation of *cry2* and *cwo* [17]. Transcripts for *tim* and *per* peak at dusk, allowing for their protein products to accumulate in the cytoplasm during the night [10, 17, 22]. A trimer of PER, TIM and CRY2 forms and migrates to the nucleus. CRY1 acts as a blue light photoreceptor which in turn degrades TIM, resulting in a PER-CRY2 dimer [23]. PER primarily serves to stabilize CRY2, which acts as an important transcriptional repressor, preventing CLK-CYC from transcribing *per* and *tim* genes, resulting in mRNA levels being minimum at dawn [23, 24]. Additionally, CWO during the night acts to repress *per* and *tim* transcription through competitive inhibition, binding to *per* and *tim* E-box promoters, preventing CLK-CYC inducing expression [17]. In the positive loop, *Clk* and *cyc* are rhythmically expressed. Transcription of *vri* and *Pdp1* occur through activation by CLK-CYC [10]. VRI acts as a transcriptional repressor and acts to delay the action of PDP1 which acts as a transcriptional activator [10, 11]. Expression of *Clk* by PDP1 occurs at dawn, causing *Clk* transcripts to peak in concentration in opposite phase to that of *per* and tim [10]. Rhythmic regulation of *cyc* transcription was observed to occur in a similar manner with E75 acting as a transcriptional repressor which delays expression of *cyc* by HR3 [16,17].

**Figure 2:**
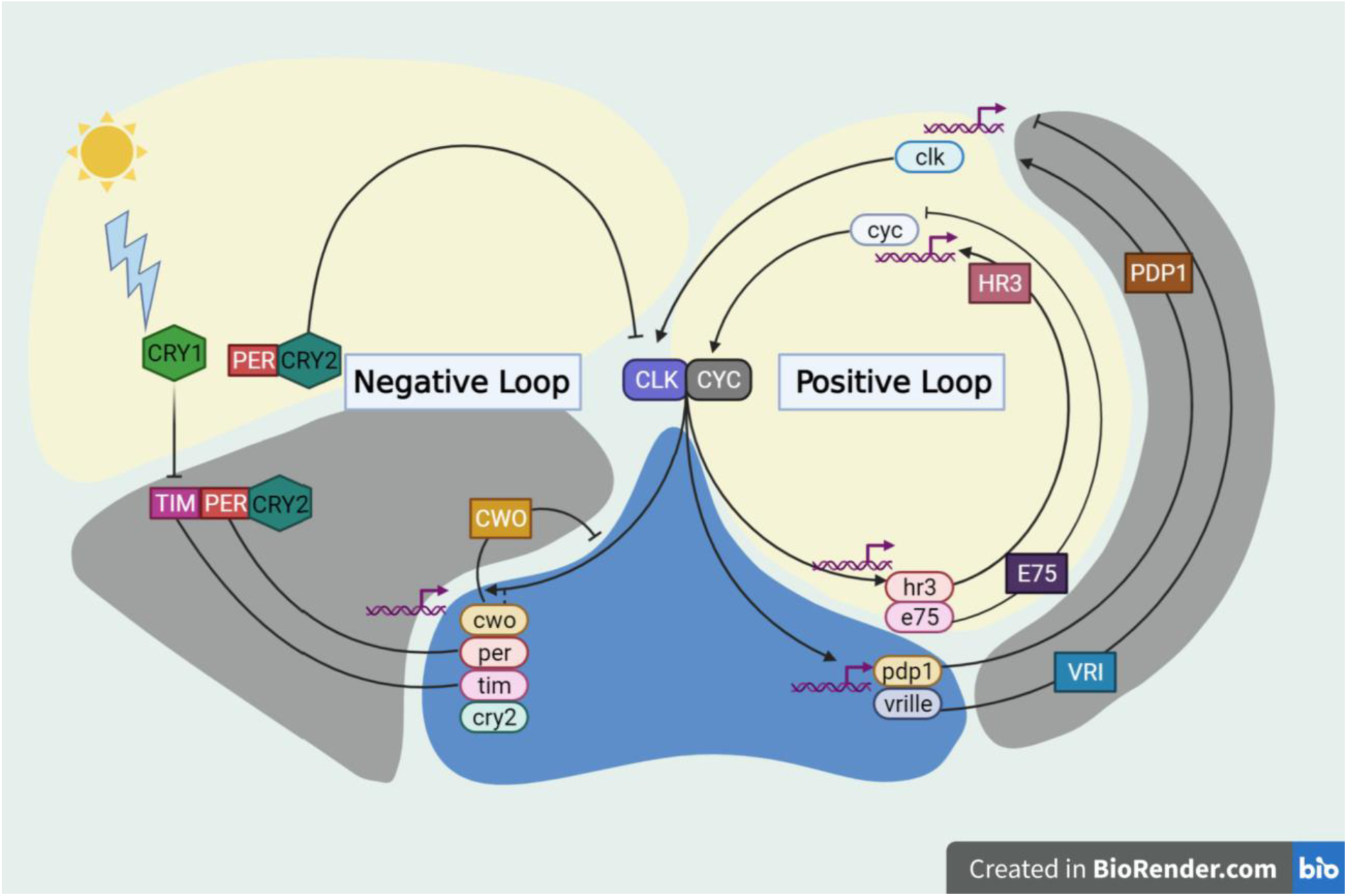
Theoretical model of circadian rhythm in *D. citri* based on curated genes and based upon circadian studies in other insect species. Arrows indicate transcriptional activation. Perpendicular lines denote transcriptional repression. Ovals represent transcripts, boxes represent proteins. The colors of the background correspond to the time of day that certain functions are (yellow = day, grey = night, blue = dusk).

## Methods

Through community curation, the *D. citri* (NCBI:txid121845) genes involved in the circadian rhythm pathway were manually annotated in genome version 3.0 [25-27], following a previously reported protocol (Fig. 3) [28]. A neighbor-joining phylogenetic tree (Fig. 4) of CRY1 and CRY2 amino acid sequences was generated with MEGA 7 (MEGA software, RRID:SCR_000667) [29] using MUSCLE (RRID:SCR_011812) for multiple sequence alignments of orthologous sequences. The circadian rhythm pathway image (Fig. 2) was created using BioRender (RRID:SCR_018361).

**Figure 3.**
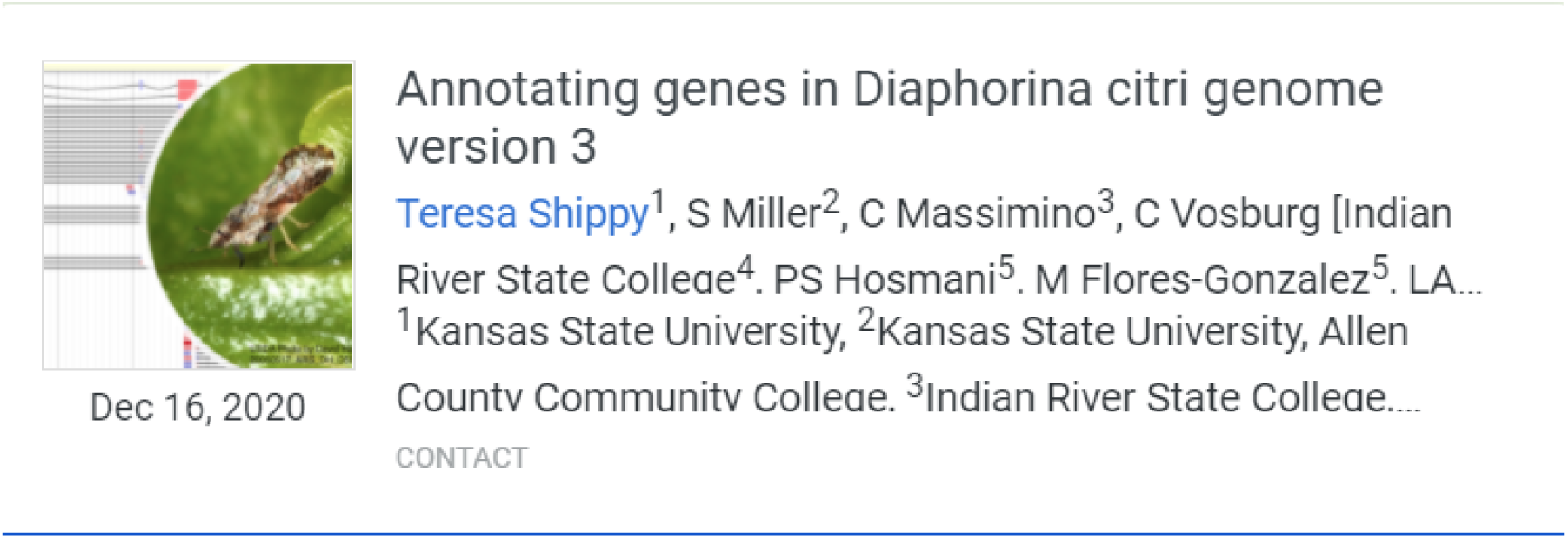
Protocol for *D*.*citri* genome community curation. [28]

**Figure 4.**
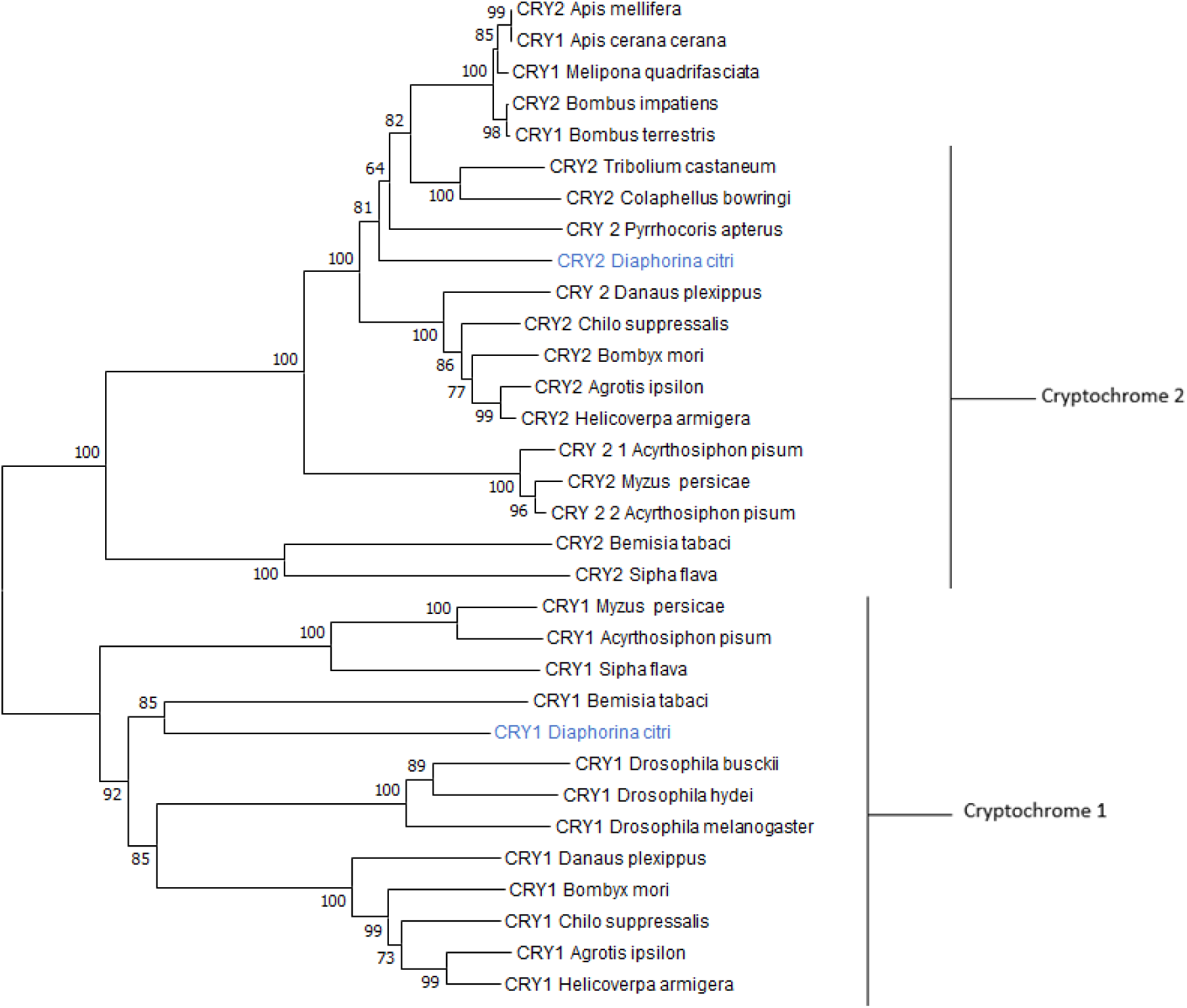
Phylogenetic tree of Cryptochrome 1 and 2 amino acid sequences. *D. citri* cryptochromes are in blue and group with their respective type of cryptochrome. *Nephila clavipes* CRY2 was used as the outgroup. The tree was made with MEGA7 using the bootstrap test (1000) replicates with the percentage of replicate trees clustered together shown next to the branches. The total number of positions used for alignment was 349 using Muscle alignment in MEGA7.

## Results and Discussion

### Core Clock Genes

Gene models were annotated in the Apollo *D. citri* v3.0 genome using the citrusgreening.org online portal [31, 32]. We identified all six of the core clock genes found in the *D. melanogaster* model [11]: *Clk, cyc, per, tim, vri* and *Pdp1*. As has been reported in *Drosophila* and the hemipteran *A. pisum*, only one copy of each gene was found in the *D. citri* genome (Table 3). However, false duplications of *Clock* and *Pdp1* resulting from genome assembly errors were found and removed from the OGS. Additionally, two of the models, *per* and *tim*, were substantially improved from the computational gene predictions. Annotation of *per* identified and corrected a splice site error, increased the amino acid length of the peptide, and improved BLAST score, query coverage and percent identity. Curation of *tim* also led to an amino acid increase from 1093 to 1109 and saw notable improvements in BLAST score, query coverage and percent identity. Notably, the added amino acids were part of a Timeless serine rich domain in the 260–292 amino acid range of the peptide, which has been identified as an important site for phosphorylation [33]. Annotation of the *vri* model may prove particularly important since lack of VRI function leads to embryonic mortality in *D. melanogaster* (Table 1) [34]. This may make *vri* a useful target for trying to control *D. citri* populations.

**Table 3:**
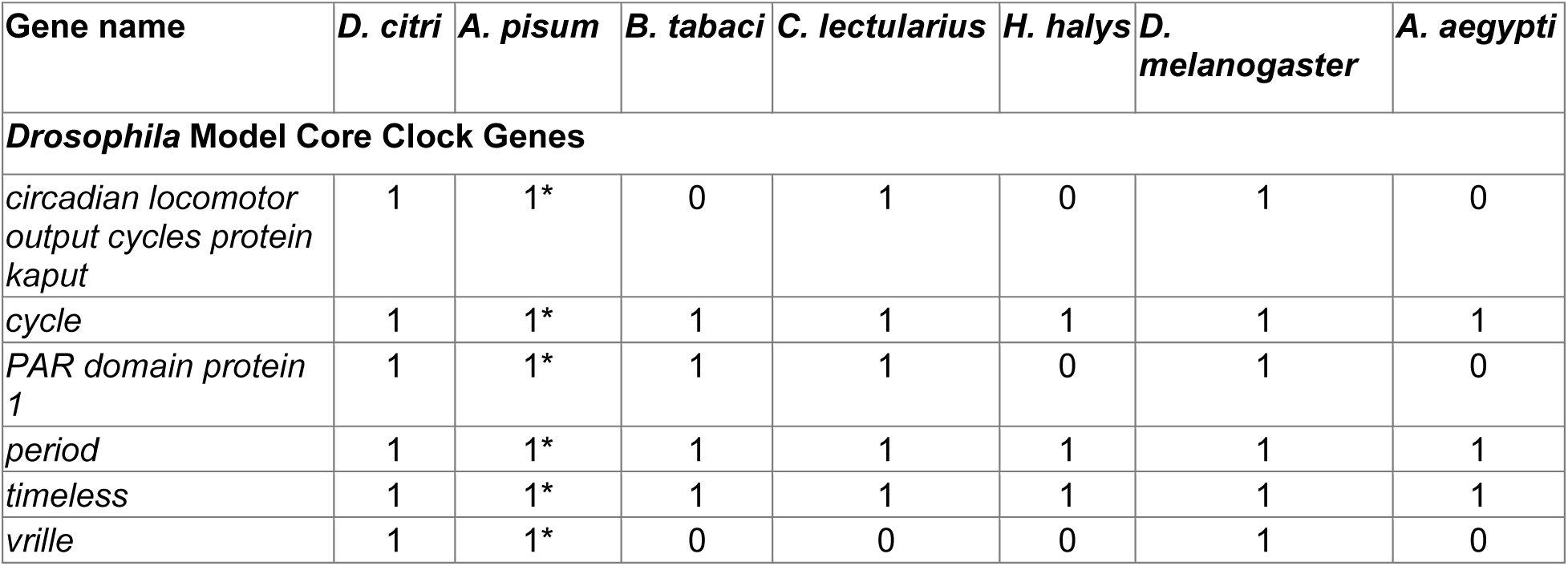
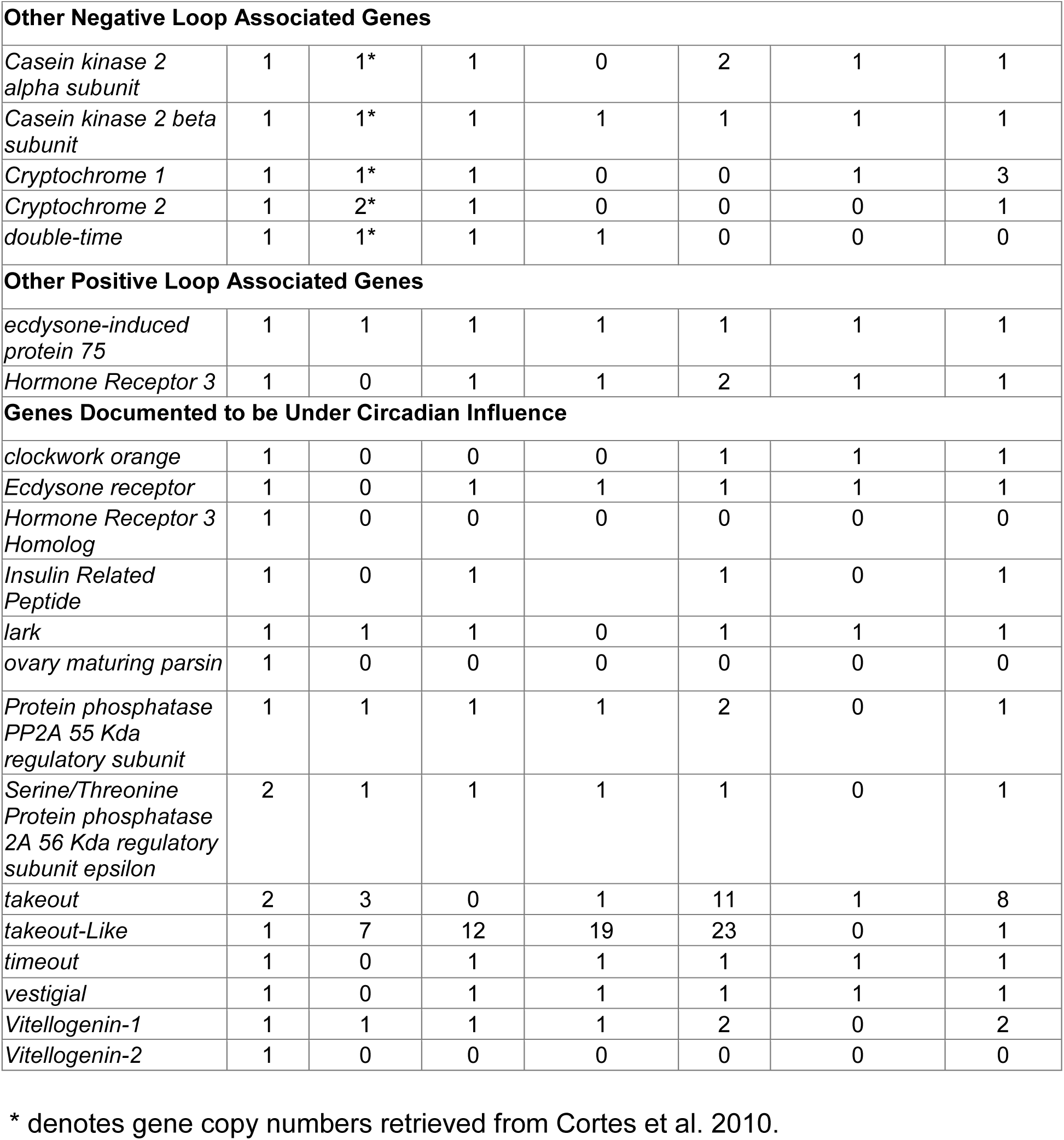
Gene copy numbers for *D. citri* compared to other insects. Numbers were obtained from NCBI unless otherwise indicated, with the exception of *D. melanogaster*, whose copy numbers were obtained from flybase.org [30].

### Other Negative Loop Associated Genes

Genes related to the negative feedback loop aside from the core clock genes include: *cry1, cry2, Casein kinase 2 alpha (Ck2α) subunit, Casein kinase 2 beta (Ck2β) subunit*, and *double-time (dbt)* (Table 3). Cryptochrome copy number variation (Fig 1) is of great importance since discrepancies between insects gives possible insight into the evolution and function of circadian rhythm in insects [14]. In *D. citri*, both *cry1* and *cry2* genes were found. A pairwise alignment between the two annotated models resulted in 41% amino acid identity with 93% query coverage. Both of the models have the DNA_photolyase (pfam00875) and FAD_binding_7 (pfam03441) domains typically found in CRY proteins. The DNA_photolyase domain is the binding site for a light harvesting cofactor, and the FAD_binding_7 domain binds an FAD molecule [35]. The BLAST results confirm that the two models were not duplications, and phylogenetic analysis was able to discern the identity of each cryptochrome (Fig. 4). The presence of *cry1* and *cry2* genes in *D. citri* reinforces the data that non-drosophilid insects possess *cry2* [14]. The presence of both genes also suggests that *D. citri* follows an ancestral butterfly-like clockwork model (Fig. 1) [14]. *D. citri* only has one *cry2* gene, while *A. pisum* has two *cry2* genes; this supports the assertion that *A. pisum* core clock genes may be evolving at a faster rate than in other hemipterans [10].

Another gene that plays an important role in the negative loop is *dbt*, also known as *discs overgrown* (*dco*). This gene encodes a circadian rhythm regulatory protein that affects the stability of PER [36] and was also annotated. The mammalian homolog of *dbt* is *Casein kinase 1* (*Ck1*) *epsilon* [37]. The *Homo sapiens* (ABM64212.1) and *D. melanogaster* (NP_733414.1) homologs both have the STKc_CK1*_*delta_epsilon (accession number: cd14125) domain at the same position, amino acids 8-282. The *D. citri* model reflects the conservation between species by also exhibiting a CK1_delta_epsilon domain located at the exact same position.

### Other Positive Loop Associated Genes

Two genes present in D. citri, *Hr3* and *E75*, are responsible for positive loop transcriptional regulation of *cyc* in *T. domestica* and *G. bimaculatus* [16,17] and a similar regulation mechanism may exist in D. citri (Fig. 2). Curation of these genes led to minimal changes to the *Hr3* gene model; however, changes in *E75* resulted in its translated peptide sequence being extended from 664 to 729. This improved its score, percent identity and query coverage when BLASTed to orthologous insects.

### Genes Documented to be Under Circadian Influence

Fourteen additional genes involved in the circadian rhythm, but not directly classified in the two feedback loops, were also annotated (Table 3). Of these genes, *takeout* is of interest as potentially a good molecular target to control *D. citri. Takeout* in *D. melanogaster* has been shown to regulate starvation response, and if the gene or protein loses function, the organism dies faster in starvation conditions when compared to the wild type (Table 1) [38]. *Takeout* and *takeout-like* in *D. citri* have low gene copy numbers when compared to other non-drosophilid insects (Table 3). The discrepancy may be due to the fact that these genes are computationally predicted in the other insects, and many sequences are short and might represent fragments of larger genes.

Another of these ancillary genes annotated was *tim* paralog *timeout*, also known as *timeless 2* [39]. *Timeout* has a number of functions and does not fit categorically into either of the feedback loops, though it is very similar structurally to *tim* [40]. To confirm that the annotated gene models were not the result of a false duplication in the genome assembly, they were compared to orthologs from other insects. The conserved domains in protein sequences for *tim* and *timeout* from *D. melanogaster* (NP_722914.3 and NP_524341.3) and *A. pisum* (ARM65416.1 and XP_016662949.1) were compared to the *D. citri* sequences. For the TIM proteins, the main conserved domain was the TIMELESS (pfam04821) domain located near the N-terminus. TIMEOUT proteins also have this N-terminal TIMELESS domain but additionally have a C-terminal TIMELESS_C (pfam05029) domain. These conserved domains were also identified in the corresponding *D. citri* protein sequences, indicating that the *timeout* model is not an artifactual duplication of *tim*. In genome version 3.0, a problem still exists causing the *timeout* sequence to be split into three different models (Table 2), despite the comparative sequence analysis that supports the presence of this gene in the genome.

## Conclusion

The circadian rhythm is an important pathway for an organism’s ability to regulate their biological systems in conjunction with the external environment [5]. A total of twenty-seven putative circadian rhythm-associated genes have been annotated in the *D. citri* genome (Table 2). This data has been used to construct models of both the positive and negative feedback loops. RNA-Seq experimental analysis over a twenty-four hour day/night cycle would provide a more definitive list of circadian rhythm associated genes in *D. citri*. Annotation of these genes provides a better understanding of the circadian pathway in *D. citri* and adds to the evidence supporting how hemipteran circadian rhythm pathways function.

## Data Validation and Quality Control

Core and auxiliary circadian rhythm genes were manually curated within the *D. citri* v3 genome. Table 2 displays the evidence utilized in these annotations as well as the completeness of the gene models. Completeness refers to the wholeness of the coding regions of these models and was determined through comparison with orthologous proteins. Curated genes were verified as orthologous to other proteins through reciprocal blasts and phylogenetic analysis.

## Reuse potential

Manual curation of these circadian rhythm genes was conducted as part of the *D. citri* collaborative community annotation project [32, 41]. These models will be incorporated into the third OGS of the Citrus Greening Expression Network (CGEN) [32]. This publicly available tool is useful for comparative expression profiling through its transcriptome data for different *D. citri* tissues and life stages. There is significant practical application of these curated models in controlling the spread of Huanglongbing. Understanding the circadian pathway and essential genes within provide opportunity for pest control strategies through implementation of molecular based therapeutics. Availability of accurate gene models brought about through our manual annotations can facilitate the design of future experiments aimed at developing CRISPR and RNAi systems to target and disrupt critical *D. citri* biological processes.

## Data Availability

The *D. citri* genome assembly, official gene sets, and transcriptome data are accessible on the Citrus Greening website [42]. The gene models will also be part of an updated OGS version 3 for *D. citri* [26, 31, 43]; the data is also available through NCBI (BioProject: PRJNA29447). All additional data supporting this article is available via the GigaScience GigaDB repository.

## Declarations

## List of abbreviations

*A. aegypti*: *Aedes aegypti*
*A. mellifera*: *Apis mellifera*
*A. pisum*: Acyrthosiphon pisum
*B. tabaci*: *Bemisia tabaci*
BLAST: Basic local alignment search tool
BLASTn: BLAST nucleotide
BLASTX: BLAST translated nucleotide
BLASTp: BLAST protein
*C. lectularius*: *Cimex lectularius*
CGEN: Citrus greening expression network
*Ck1*: *Casein kinase 1*
*clk*: *Clock*
*CLOCK*: *Circadian locomotor output cycles protein kaput*
CLas: Candidatus Liberibacter asiaticus
*cry1*: *Cryptochrome 1*
*cry2*: *Cryptochrome 2*
*cwo*: *clockwork orange*
*cyc*: *cycle*
D. citri: *Diaphorina citri*
*D. melanogaster*: *Drosophila melanogaster*
*D. plexippus*: *Danaus plexippus*
*dbt*: *double-time*
*dco*: *discs overgrown*
DNA: Deoxyribonucleic Acid
*E75*: *ecdysone-induced protein 75*
FAD: Flavin adenine dinucleotide
FASTA: FAST-All
GigaDB: GigaScience DataBase
*H. halys*: *Halyomorpha halys*
HLB: Huanglongbing
*Hr3*: *Hormone receptor 3*
Iso-Seq: Full-length isoform sequencing
MCOT: MAKER Cufflinks Oases Trinity
*Mip*: *myoinhibitory peptide*
NCBI: National center for biotechnology information
OGS: Official gene set
*PDFR*: Pigment-dispersing factor receptor
*Pdp1*: *PAR domain protein 1*
*per*: *period*
*PP2A*: *Protein phosphatase 2A*
RNA-Seq: RNA-sequencing
RNAi: RNA interference
STKc: Serine/threonine kinase catalytic domain
*T. castaneum*: *Tribolium castaneum*
*tim*: *timeless*
*vri*: *vrille*

## Ethical Approval

Not applicable.

## Consent for publication

Not applicable.

## Authors’ contributions

SJB, WBH, TD and LAM conceptualized the study; TD, SS, TDS, CM, CV and SJB supervised the study; TD, SS, LAM and SJB contributed to project administration; LMD, CM, JN, YO, ME, ND, RM, AN, RH, KG, MK, MTH conducted the investigation; PH, SS, MF-G contributed to software development; SS, TDS, PH and MF-G developed methodology; SJB, WBH, TD and LAM acquired funding; MR, LMD, TP and YO prepared and wrote the original draft; LMD, TP, WBH, TDS, JB, YO, ME, RM, AN, reviewed and edited the draft.

## Acknowledgements

We would like to thank Helen Wiersma-Koch (Indian River State College), Elizabeth Michels, Sarah Michels, and Tanner Wise for assistance

